# Linking Host Covariates to COVID-19 Vaccine-Induced Antibody Dynamics in a Large Healthcare Worker Cohort

**DOI:** 10.64898/2026.06.11.731600

**Authors:** Clemens Peiter, Emma Sala, Giuseppe De Palma, Carlotta Zunarelli, Michelle B. Hanson, Mahsa Abedini, Evelina Tacconelli, Paolo Boffetta, Jan Hasenauer

## Abstract

Antibody responses to COVID-19 vaccination vary widely across individuals, but the factors shaping these differences and their relation to underlying host-response processes are still not fully understood. Here, we combined longitudinal serological data from 4,553 healthcare workers with a nonlinear mixed-effects model to quantify determinants of vaccine-induced antibody dynamics. This framework captured both population-level trends and inter-individual variability in anti-spike antibody trajectories after vaccination. We found that age, sex, and pre-vaccine infection significantly influenced response dynamics. Male sex and increasing age were associated with weaker initial antibody production but slower waning, while individuals with a pre-vaccine infection had pre-existing anti-spike antibodies and slower initial antibody production and faster waning. These effects translated into substantial long-term differences in predicted antibody levels, reaching up to a 27% decrease in antibody levels one year after the third vaccination. Our results provide a dynamic view of the determinants of vaccine-induced humoral immunity and establish a framework for analyzing heterogeneous immune responses in large longitudinal cohorts.

## Introduction

In late 2019, coronavirus disease 2019 (COVID-19), caused by severe acute respiratory syndrome coronavirus 2 (SARS-CoV-2), emerged as a global health crisis and has since resulted in millions of deaths worldwide [1]. Although population immunity acquired through infection and vaccination has substantially reduced the burden of severe disease, SARS-CoV-2 continues to circulate, and newly emerging variants remain capable of causing large numbers of infections [2]. Vaccination has therefore remained a central component of public health strategies to limit transmission, reduce hospitalizations and mortality, and protect vulnerable populations. Vaccines were rapidly developed and deployed at unprecedented scale [3,4], and proved highly effective in reducing severe COVID-19 and COVID-19–related mortality [5,6]. However, we and others observed that the magnitude and durability of vaccine-induced immune responses vary considerably between individuals [7], with important implications for the duration of protection and the design of vaccination strategies [8].

Computational modeling has played a key role throughout the pandemic, supporting quantitative analysis across multiple biological and epidemiological scales. These efforts have ranged from population-level models of disease spread [9], over within-host models of viral dynamics [10], to mathematical descriptions of immune responses following vaccination [7,11–13]. Observational studies have identified host-related and clinical covariates associated with variation in post-vaccination antibody levels, including age, sex and prior infection status [7,11,14]. Such studies have provided important evidence for heterogeneity in vaccine responses, but they have predominantly relied on cross-sectional analyses or regression-based approaches applied to isolated time points. As a result, they are well suited to identifying associations with antibody magnitude at specific times, but less suited to explaining how covariates shape the temporal evolution of the response.

This distinction is important because vaccine-induced immunity is inherently dynamic. Antibody levels are determined not only by peak response magnitude, but also by the rates of induction, amplification and decline, and different factors may act on different components of this process [8]. Distinguishing these effects is essential for understanding why individuals with similar early responses may diverge substantially at later time points, and why comparable antibody levels measured at a single visit may reflect different underlying response trajectories. A dynamic modelling framework can therefore provide insights that are inaccessible to cross-sectional analyses, by linking observed longitudinal data to interpretable features of the host response. Such information is relevant for the rational timing of booster vaccinations, for the identification of groups with less durable immunity, and more broadly for understanding how host factors modulate vaccine responsiveness [15].

Despite this need, dynamic modeling studies of COVID-19 have focused primarily on viral kinetics and host–pathogen interactions during infection [16–18]. Mathematical models of vaccine-induced antibody dynamics have been developed [12,13], but their application has largely centered on describing average trajectories rather than quantifying how host-related factors shape the underlying response processes in large heterogeneous populations. Furthermore, the datasets used for dynamic modeling studies were generally small, including at most a few hundred individuals. Consequently, the relationship between known covariates of vaccine responsiveness and the dynamic features of the antibody response remains insufficiently resolved.

Here, we address this gap by modeling longitudinal serological data from the ORCHESTRA cohort [11] with a nonlinear mixed-effects (NLME) model of vaccine-induced antibody dynamics. The NLME framework allows us to jointly characterize population-level response patterns and inter-individual variability, while relating host covariates to interpretable features of antibody production and waning. We analyzed 4,553 healthcare workers from the Bologna and Brescia study sites of the ORCHESTRA cohort, for whom repeated anti-spike antibody measurements and vaccination records were available across at least four time points. After establishing model applicability and practical identifiability of the parameters, we used the calibrated model to quantify the effects of age, sex and pre-vaccine infection on antibody dynamics. We show that these covariates systematically reshape antibody trajectories and can give rise to substantial long-term differences in predicted antibody levels. Together, these results identify distinct effects of age, sex, and prior infection on the immunological processes controlling the magnitude and persistence of vaccine-induced antibody responses and demonstrate how NLME modeling can be used to quantify immune heterogeneity in large longitudinal cohorts.

## Material and methods

### Data collection and processing

Data were collected within the “Connecting European Cohorts to Increase Common and Effective Response to SARS-CoV-2 Pandemic” (ORCHESTRA) project and have been described previously by Leomanni et al. [11]. In brief, ORCHESTRA includes, amongst other participant groups, more than 60,000 healthcare workers from several European hospitals. For this study, we were provided with a subset of healthcare workers from Bologna and Brescia, Italy. These sites contained longitudinal data with repeated antibody measurements, vaccination records and follow-up across multiple visits. Blood samples were used to quantify binding antibodies (BAU) against the SARS-CoV-2 spike protein (anti-S antibodies), and covariate information including age, sex, job, vaccination status, and pre-vaccine infection was available for each participant. To ensure sufficient information for individual-level parameter estimation, we further restricted the analysis to participants with samples from at least four time points and applied additional basic filtering criteria to arrive at our analysis set of 4,553 individuals (Fig 1A). In particular, individuals with evidence of infection during the observation period were excluded since the time of infection was not known and could therefore not be incorporated reliably into the modeling framework. Details of all exclusion criteria are provided in the Supporting information <u>S1 Text</u>, section *Filtering steps applied to the dataset to arrive at the analysis set*.

**Figure 1.**
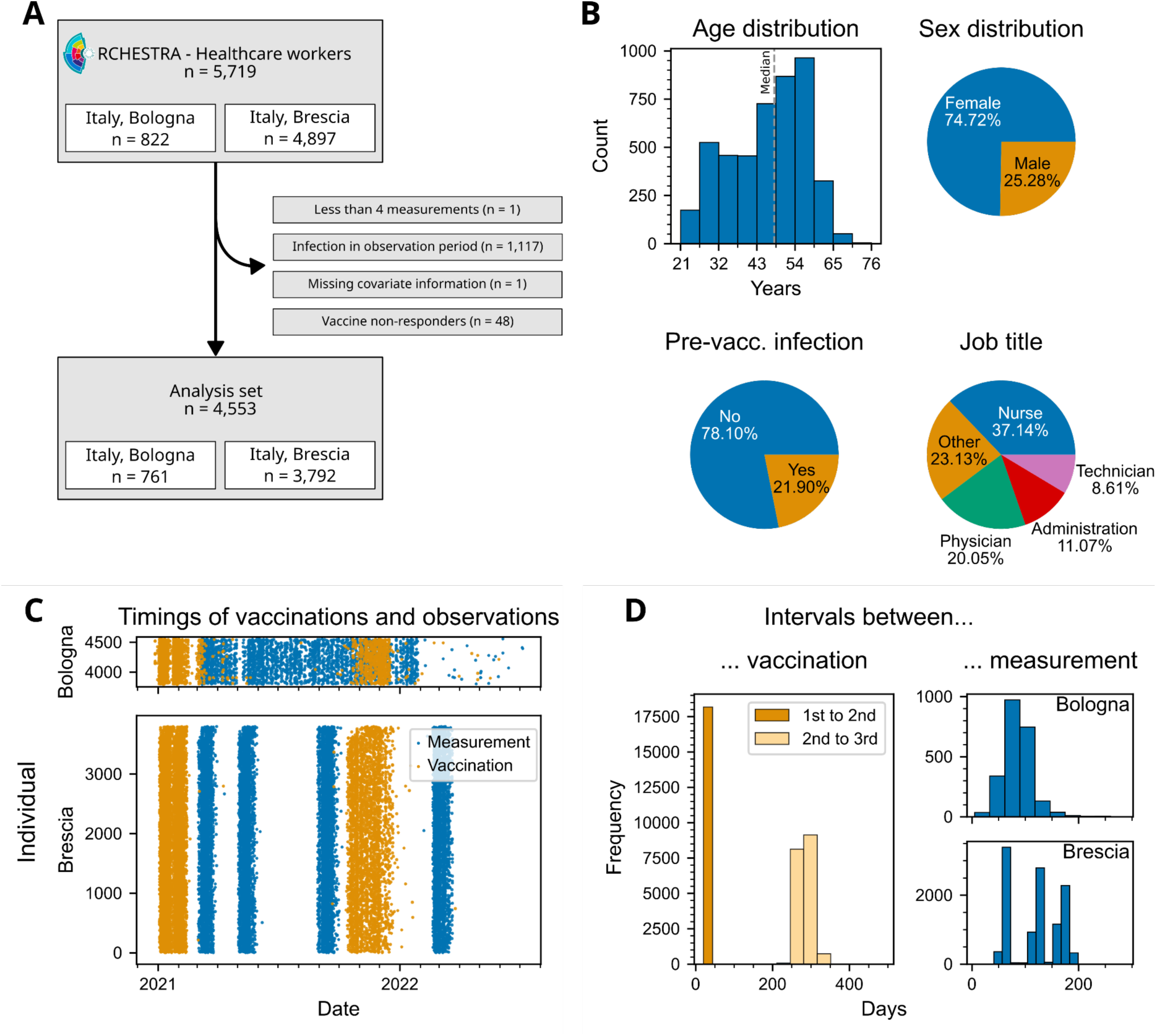
Overview of the (filtered) ORCHESTRA healthcare worker dataset for Bologna and Brescia. (A) Flow diagram showing total number of individuals and applied filtering steps to arrive at the analysis set. (B) Histogram and pie charts showing the distribution of available covariates in the analysis set. (C) Scatter plot displaying the measurement and vaccination time points for each individual. (D) Histograms of time intervals between vaccination and measurements for all individuals in the analysis set.

### Individual-level model of antibody dynamics

To describe vaccine-induced antibody dynamics, we used ordinary differential equations (ODEs), a well-established framework for mechanistic modeling of dynamic biological processes [19]. Following vaccination, host cells transiently express the vaccine-encoded antigen, which triggers activation of the humoral immune response and induces antibody production [20]. As antigen expression wanes, antibody levels gradually decline. Although the underlying response involves multiple cellular compartments and regulatory processes, the available data consist only of longitudinal antibody measurements and vaccination times. We therefore considered a coarse-grained mechanistic model that captures the effective induction and decline of antibodies following vaccination.

As a starting point, we considered a previously published ODE model of post-vaccination antibody dynamics [13]. In our setting, however, this formulation proved practically non-identifiable on the individual level and produced biased parameter estimates on the population level (<u>S1-S3 Figs</u>; see also <u>S1 Text</u>, section *In silico testing of the modeling and parameter estimation pipeline*). We therefore simplified the model further by replacing the original logistic formulation of intrinsic antibody dynamics with first-order exponential decay (see <u>S1 Text</u>, section *Comparison of model from dePillis et al. (2023) and our model*). The resulting model is

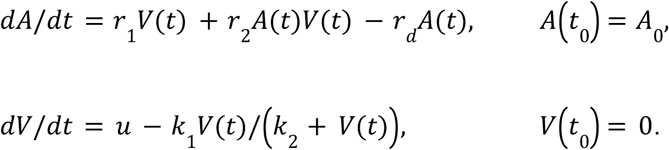

Here, *A(t)* denotes the latent antibody level and *V(t)* an unobserved state variable representing a proxy for vaccine-transfected, antigen-expressing cells at time *t*. The input *u* denotes a normalized vaccine input function that is equal to 1 on the day of vaccination and 0 otherwise. Following vaccination, *u* increases *V*, while *V* decreases through a saturable loss term parameterized by *k_1_* and *k_2_*. Antibodies are produced at the effective rate *r_1_V(t)*, which is proportional to the amount of antigen-expressing cells. The additional term *r_2_A(t)V(t)* represents an amplification of antibody production that depends jointly on the current antibody level and antigen availability, thereby capturing memory-like effects in a coarse-grained manner. In the absence of antigen expression, that is, for *V=0*, the model reduces to *dA/dt = -r_d_A(t)*, so that antibody levels decay exponentially with first-order decay rate parameter *r_d_*. The initial antibody level *A_0_* accounts for pre-existing antibodies before the first vaccination; for individuals without pre-vaccine infection, *A_0_*was fixed to zero. A schematic representation of the model is shown in Fig 2A.

**Figure 2.**
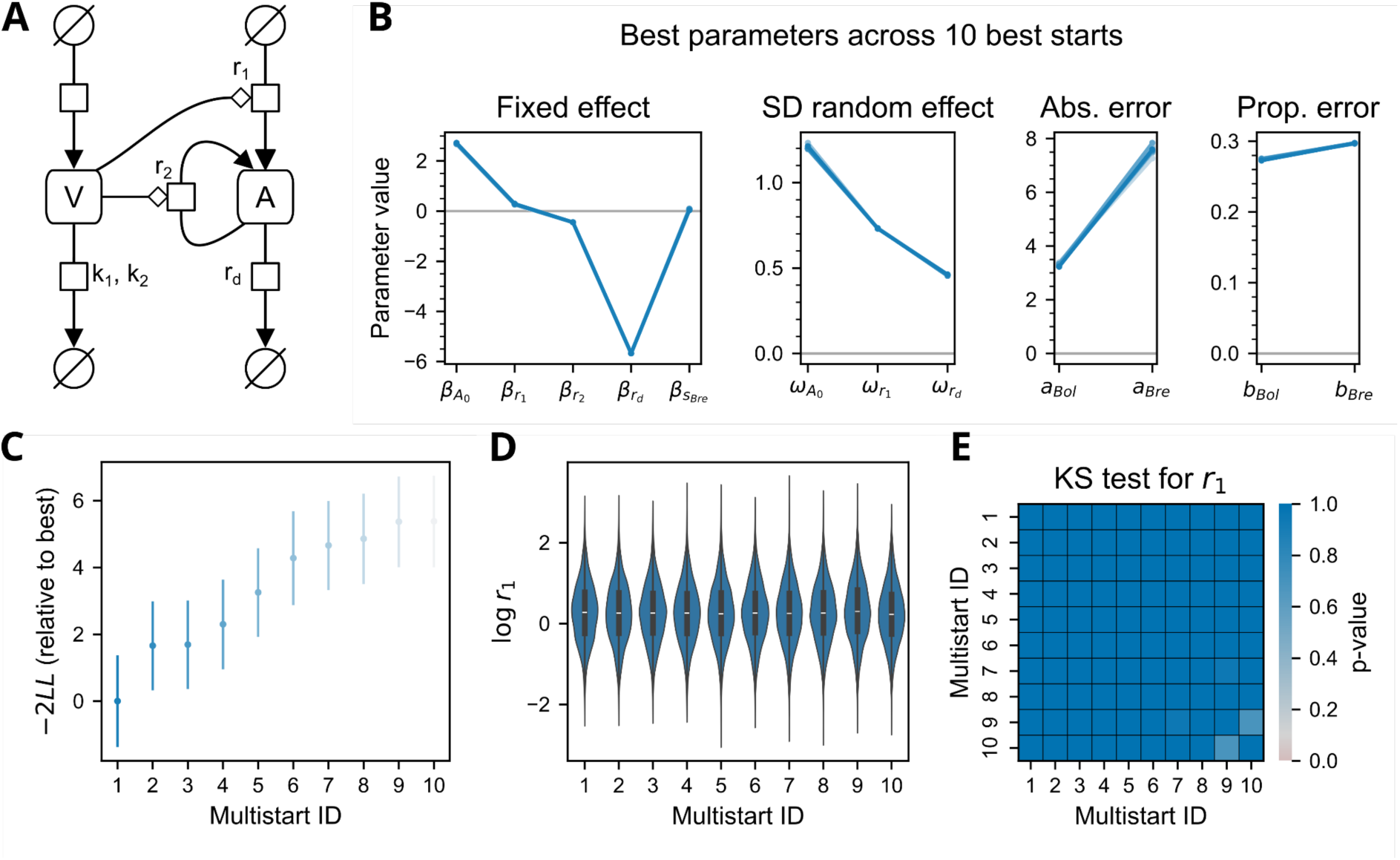
Summary of multistart optimization result for the simplified model. (A) Schematic representation of the ODE model in SBGN format. (B) Graph showing the ten best parameter estimates from our multistart approach according to likelihood. (C) Estimated likelihoods and standard errors of the likelihood for the ten best starts. The likelihood value is relative to the best estimated likelihood. (B,C) Lighter colors correspond to worse likelihood values. (D) Violin plots showing samples from the estimated random effects distribution for parameter *r_1_*for the ten best starts. (E) Heatmap of Kolmogorov-Smirnov test results, calculated for the samples and starts in (D).

To connect the latent model state to the measured antibody concentrations, we used an observation model that accounts for relative and absolute measurement noise, censoring and general differences between the antibody measurement techniques employed by the different cohorts. For an individual *i* from cohort *c* ∈ {*Bol, Bre*}, where *Bol* and *Bre* denote the Bologna and Brescia cohorts, respectively, the measurement model was

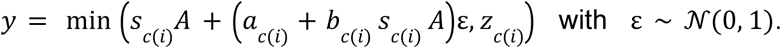

Here, *s_c(i)_*denotes the scaling factor, *a_c(i)_*and *b_c(i)_*denote the additive and proportional error parameters, and *z_c(i)_* denotes censoring level. The subscript *c(i)* encodes to which cohort individual *i* belongs. The observation model defines the conditional probability of observing a measurement *y* given the state of an individual: (i) the likelihood contribution of an uncensored measurement (*y* < *z_c(i)_*) is 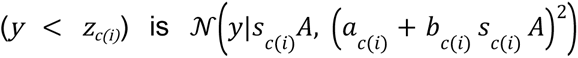 , while (ii) the likelihood contribution of a censored measurement (*y = z_c(i)_*) is given by the integral

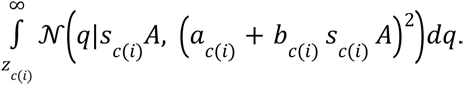

To avoid structural non-identifiability, the scaling parameter for the Bologna cohort *s_Bol_*was fixed to 1, whereas the corresponding parameter for Brescia *s_Bre_* was estimated.

For numerical robustness and to ensure positivity of all ODE model parameters, we performed estimation on the log scale. Error model parameters were estimated on a linear scale and are also restricted to be positive. Accordingly, the individual-level parameter vector was defined as

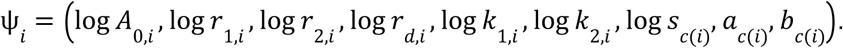

### Population-level description of vaccination response

To assess the drivers of heterogeneity in the vaccination response, we embedded the individual-level model in an NLME framework [21]. This approach allows us to describe both the population-average antibody dynamics and the inter-individual variability around this average.

At the population level, only a subset of parameters was allowed to vary between individuals. Specifically, *A_0_*, *r_1_*, and *r_d_* were subject to inter-individual variability and were therefore modelled using random effects. In contrast, *r_2_*, *k_1_*, and *k_2_* were treated as fixed across individuals as previous analyses with inter-individual variability included revealed that it provided no benefit to the estimation with our dataset (<u>S4 Fig</u>). In addition, *k_1_* and *k_2_* were fixed to previously reported and biologically reasonable values [13] (see <u>S1 Table</u>).

For parameters subject to inter-individual variability, we assumed a log-normal distribution, meaning that for θ ∈ {*A_0_, r_1_, r_d_*} it holds that

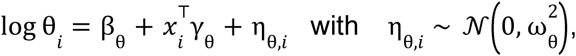

where β_θ_is the population-level intercept, *x_i_*is the covariate vector for individual *i*, γ_θ_the corresponding covariate effects, η_θ,*i*_ the individual-specific random effect, and ω_θ_ the standard deviation of the random effect. We denote as the baseline model, where all covariate effects were set to zero, that is, γ_θ_ *=* 0. Random effects were assumed to be independent, resulting in a diagonal covariance matrix. For consistency, visualization purposes, and ensuring that parameters stay positive we also write log θ = β_θ_ for θ ∈ {*r_2_, s_Bre_*}.

### Parameter estimation

Population-level parameters were estimated using a stochastic approximation expectation-maximization (SAEM) algorithm [22], considering data from Bologna and Brescia. In the baseline model, the unknown parameters comprised the population-level intercepts for the structural parameters, the variances of the random effects for *A_0_*, *r_1_*, and *r_d_*, and the cohort-specific observation parameters for scaling and measurement noise. Specifically, we estimated the fixed effects β_θ_ for all structural parameters that were not fixed a priori, the random-effect standard deviations ω_θ_for θ ∈ {*A_0_, r_1_, r_d_*}, and the additive and proportional error parameters for both cohorts.

We combined SAEM with a multistart strategy to ensure the reproducibility of parameter estimates. For the multistart procedure, initial parameter values for *β_θ_*were sampled from uniform distributions where the bounds were chosen based on the data range (<u>S1 Table</u>), which should cover all biologically plausible values. Each sampled initialization was used in one independent optimization run, which we refer to as a single start. We performed 50 starts per model or more if the estimation had not converged.

After estimation of the population-level parameters, we sampled multiple individual-specific parameters per individual from the conditional distribution of the random effects given the observations and the estimated population parameters. These samples were used for simulation and covariate analysis. In addition, empirical Bayes estimates (EBEs) were obtained for each individual from the corresponding conditional distribution and used to summarize individual-level fits.

### Analysis of fitting results

To assess the fit quality at the individual level, we simulated the model using the EBE for each individual and compared the resulting trajectory with the observed data using a residual-based root mean squared error (RMSE). In case of censored measurements, the RMSE was zero if both the simulation and measurement were in the censoring interval. Otherwise the RMSE corresponded to the distance between the two points. We further estimated the log-likelihood and parameter standard errors by importance sampling conditional on the fitted model parameters. Parameter estimates are reported with the standard errors or corresponding 95% confidence intervals.

To assess whether the results of the different good optimization runs within the multi-start are comparable, we used the nonparametric approach proposed by Cassidy et al. [23]. Here, we draw as many samples from the estimated distributions of the random effects as there are individuals in the dataset. We do this for each of the best starts and perform a Kolmogorov-Smirnov (KS) test on the samples to check whether the samples come from the same distribution. A small p-value indicates that the estimated distributions differ from one another. Depending on the number of tests we apply multiple testing corrections using the Bonferroni method. For parameters without random effects, convergence was evaluated by comparing the estimates across starts relative to their estimated standard errors.

### Estimation of covariate effects

To identify covariates associated with inter-individual differences in antibody dynamics, we followed an iterative procedure similar to Ayral et al. [24].

At each iteration, we drew 50 samples per individual from the conditional distribution of the random effects given the data and the fitted model starting with the baseline model. For the continuous covariate age, associations with random effects were assessed using Pearson’s correlation coefficient. For categorical covariates, differences between groups were assessed using analysis of variance (ANOVA). Covariates showing significant associations (significance level 0.05 with multiple testing correction using the Bonferroni method) with the random effects were considered as candidates for inclusion in the model.

For each candidate covariate, an NLME model was constructed using the parametrization scheme outlined in the section *<u>Population-level description of vaccination response</u>*. For example, when testing an effect of age on *r_1_*, the corresponding model term was written on the log scale as

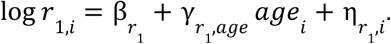

More generally, covariate effects entered through the fixed-effects part of the mixed-effects model, while the random-effects structure was retained. The candidate models were fitted using the method described in the section *<u>Parameter estimation</u>*, yielding maximum likelihood estimates Θ̂ as well as the corresponding likelihood values *L*(Θ̂). Furthermore, we obtained standard errors for the covariate-effect parameters based on which we tested – using the Wald statistics – whether the corresponding coefficients differed significantly from zero.

Model selection was used to select the best model among the candidate models, accounting for statistical significance as well as goodness of fit. A covariate effect was retained only if the corresponding parameter was significantly different from zero and the corrected Bayesian information criterion [25] improved by at least 10 as it provides strong evidence for the new model [26]. The corrected Bayesian information criterion was computed as

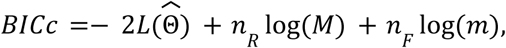

where *L*(Θ̂) is the log-likelihood of the estimated population parameters, *M* the number of individuals, *m* the total number of observations, and *n* and *n* the numbers of estimated random-effects and fixed-effects parameters, respectively.

Following this procedure, we add one new covariate to NLME models from the last iteration until no further improvement was achieved.

### Data availability

The experimental datasets of this study can be made available in de-identified format upon reasonable request to the principal investigators of the participating cohorts (Paolo Boffetta for Bologna and Giuseppe De Palma for Brescia). Additionally, we created a synthetic dataset that can be used in our estimation and plotting routines that is published with all other code on Zenodo.

### Software

We used Monolix version 2023R1 [27] and the provided R package lixoftConnectors for parameter estimation in the NLME setting, conditional distribution sampling, likelihood estimation, and standard error estimation. Additionally, we developed our own interface in R version 4.4.2 [28] for a multistart optimization procedure that could be parallelized on high performance computing infrastructure. The analyses of parameter estimates and covariate effects were performed in Python version 3.12.3. All simulations for visualization were performed using libRoadRunner version 2.8.0 [29,30]. The plots were created in Matplotlib version 3.10.8 [31]. All code is made available on Zenodo at https://doi.org/10.5281/zenodo.20644099.

## Results

### ORCHESTRA cohort supports reproducible inference of post-vaccination antibody dynamics

To investigate determinants of post-vaccination antibody dynamics in a large heterogeneous population, we analyzed longitudinal serological data from the ORCHESTRA project. The analysis focused on participants from Bologna and Brescia, Italy, as these sites provided the most complete longitudinal records for dynamic modeling. After multiple filtering steps (Fig 1A), longitudinal data from 4,553 healthcare workers comprised the analysis set (n=761 for Bologna; n=3,792 for Brescia). The median age was 48 years, and about 75% of the cohort were female (Fig 1B). More than half of the participants had occupations with close patient contact, including nurses and physicians (57.19%), whereas the remainder worked in roles not necessarily involving close patient contact, such as administration or technical services (42.81%). In addition, a subset of individuals had experienced infection before their first vaccination (21.9%). For each participant, binding antibodies against the SARS-CoV-2 spike protein were measured at at least four visits (Fig 1C). The two cohorts differed somewhat in their sampling patterns: measurement time points from Bologna were more irregularly distributed across individuals, whereas sampling in Brescia was more regular. Vaccination intervals were broadly consistent between the two sites, with little variability between the first and second vaccination and greater variability between the second and third vaccination (Fig 1D).

To assess whether reliable parameter estimation was possible from these data, we calibrated the baseline model (Fig 2A) with minimal covariate effects on the full dataset and performed complementary *in silico* analyses (see <u>S1 Text</u>, section *In silico testing of the modeling and parameter estimation pipeline*). Parameter estimation proved reproducible across independent optimization runs (Fig 2B-E). For comparable likelihood estimates, particularly when the standard errors of the likelihood overlapped (Fig 2C), the corresponding parameter estimates were nearly identical (Fig 2B). To further assess reproducibility of the estimated population distributions, we applied the identifiability analysis proposed by Cassidy et al. [23]. An example is shown in Fig 2D and E, where the estimated distributions from the ten best optimization runs are virtually indistinguishable. The random effects sampled for each parameter across these runs (Fig 2D) appeared to originate from the same distribution, as confirmed by the KS test (Fig 2E). In addition, the baseline model showed accurate parameter recovery in the *in silico* setting (<u>S3 Fig</u>). The estimated half-life of antibodies, determined by *r_d_*, was approximately 201 days (95% CI: 195-203 days) (Fig 2B). For all subsequent fitting procedures and covariate analyses, we therefore used the baseline model and selected the fit corresponding to the highest likelihood estimate.

### Model captures antibody dynamics at the individual and population levels

To evaluate whether the baseline model not only yielded reproducible parameter estimates but also provided an adequate description of the observed antibody dynamics, we assessed its goodness of fit at both the individual and population levels. This analysis is distinct from the reproducibility and identifiability assessment above: whereas the previous section addressed whether model parameters can be estimated reliably, here we examined whether the fitted model captures the structure of the measured data sufficiently well to support downstream covariate analysis. To this end, we sampled from the conditional distribution of the individual-specific parameters given the data and simulated antibody trajectories for all individuals (Fig 3).

**Figure 3.**
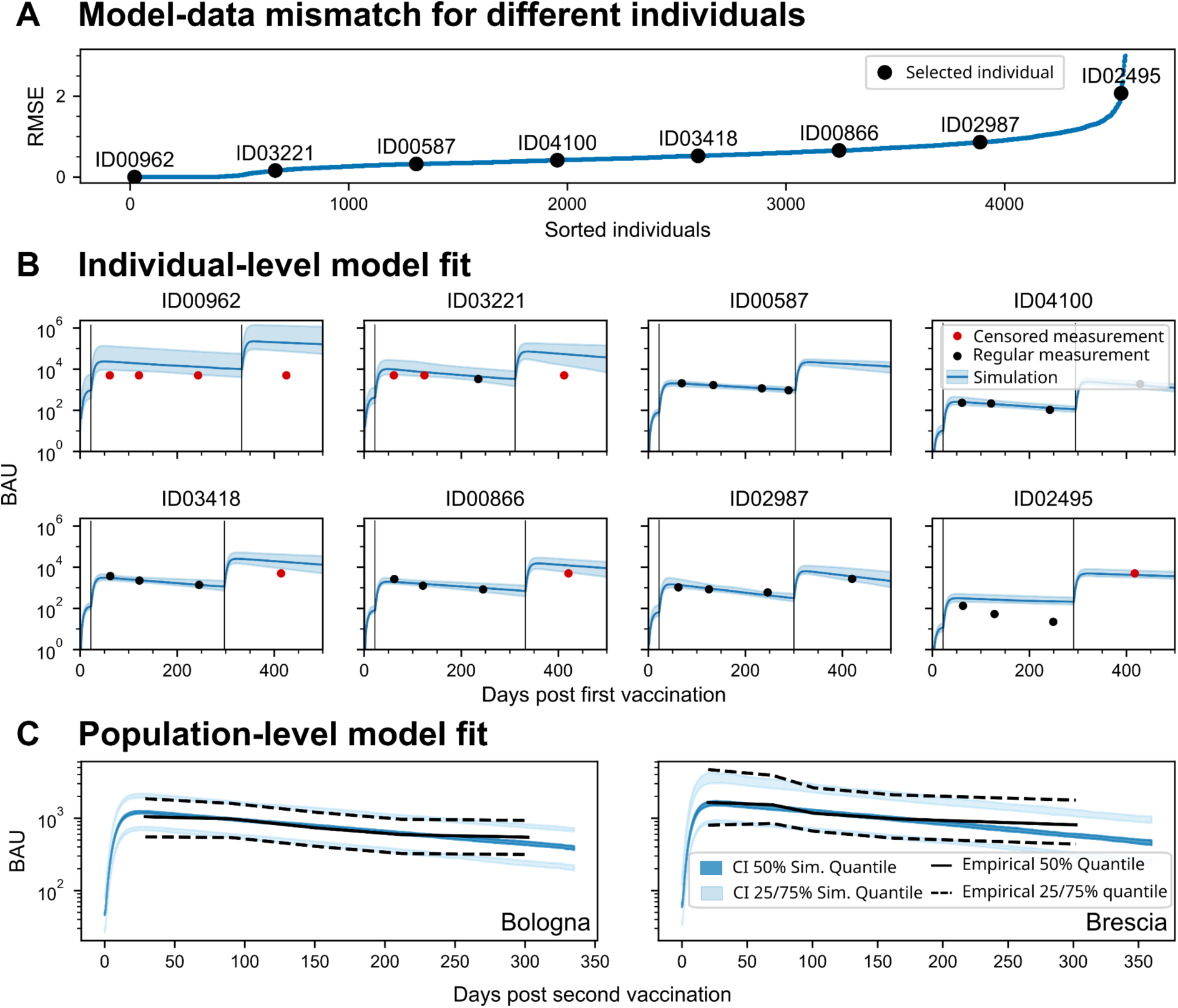
Individual and population model simulations using the optimal parameter estimates. (A) RMSE between the simulation with the EBE and the data, evaluated for all individuals. The individuals are sorted from lowest to highest RMSE. Black dots indicate selected individuals used for further visualization of the individual model fits. (B) Individual-level model fits for representative individuals with 95% credible intervals from conditional distribution samples. The vertical black lines indicate vaccination events. (C) Population model fits show a comparison between simulated quantiles and empirical quantiles for observations between second and third vaccination. The bands of the simulated quantiles display the 95% confidence interval of the quantiles. The empirical quantiles are calculated on binned measurements between second and third vaccination.

At the individual level, the model provided a good description of the observed trajectories. This was reflected in the distribution of the residual-based RMSE values (Fig 3A) and in representative individual fits (Fig 3B). In interpreting these results, it is important to note that the dataset contains censored measurements, particularly for individuals with very strong antibody responses. In particular, many individuals with strong antibody responses contributed repeated right-censored measurements, and simulations exceeding the censoring threshold were treated as consistent with these observations in the RMSE calculation. Thus, low RMSE values do not only reflect small deviations between simulated and observed values on the measurement scale, but also the fact that the model correctly captured trajectories entering the censored regime. Only for a small number of individuals with the poorest fits observations were not covered by the 95% prediction interval of the simulations. In addition, residual analysis showed no major systematic deviations and supported the adequacy of the chosen mixed error model (<u>S5 Fig</u>).

To assess the fit at the population level, we focused on the period between the second and third vaccination, where the data were most informative because both the number of measurements and the extent of observable dynamics were greatest. We evaluated the population-level fit separately for Bologna and Brescia and found good agreement of simulated and empirical quantiles derived from the data for both cohorts (Fig 3C). Together, these results indicate that the NLME model captures the observed antibody dynamics sufficiently well at both the individual and population levels and therefore provides a suitable basis for downstream analysis of covariate effects.

### Covariates shape the dynamics of vaccine-induced antibody responses

To understand the mechanistic basis of inter-individual differences in vaccine-induced antibody responses, we investigated how covariates relate to interpretable features of the model dynamics, in particular antibody production, decline and pre-existing antibody levels.

In the first screening step we analyzed associations between covariates and the individual-specific random effects inferred under the baseline model without covariate effects, and then incorporated candidate covariate effects directly into the NLME model for refitting and model selection.

This screening analysis identified several covariate–parameter associations (Fig 4A; example shown for age and pre-vaccine infection). In particular, both age and sex were associated with the random effects of *r_1_* and *r_d_*, suggesting effects on antibody production and decline. Pre-vaccine infection affected the initial antibody levels in *A_0_* and showed strong associations with *r_1_*, and *r_d_*. In addition, cohort membership was associated with variation in *A_0_*. Overall, the screening step identified a limited set of plausible covariate–parameter relationships for formal testing in the population model.

**Figure 4.**
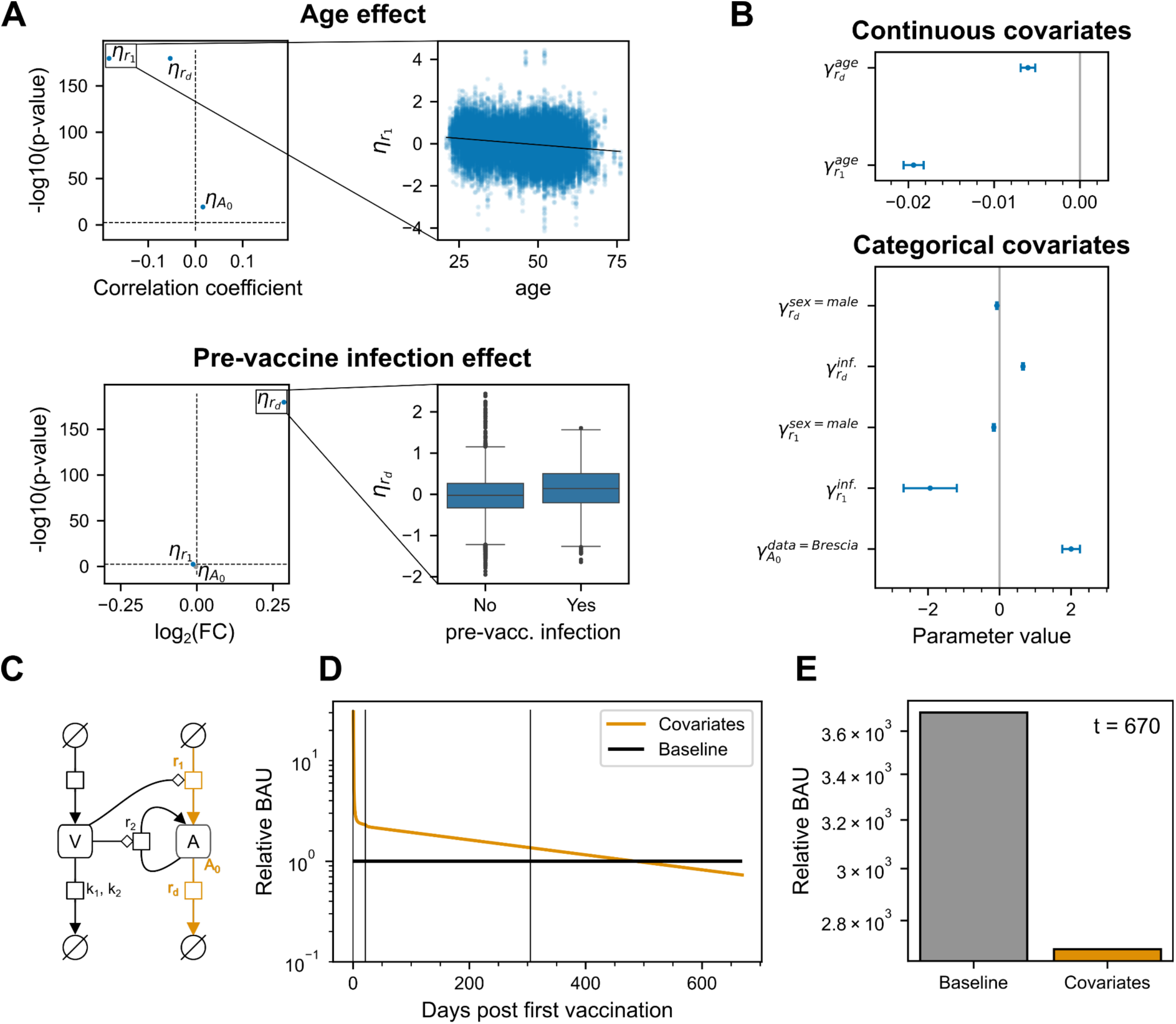
Analysis of covariate-parameter relationships. (A) Volcano plots on the left show the Pearson correlation coefficient between random effect and covariate or the mean fold-change (FC) covariates and associated p-values of the covariate effect. Scatter and boxplots on the right show examples highlighted by the black rectangles of the random effect samples plotted against the covariate. (B) All estimated covariate effects included in the final model with their estimated standard errors. The grey vertical line indicates no covariate effect. (C) Schematic overview of the ODE model with covariate-affected parameters and pathways highlighted in orange. (D) A simulation comparing the predicted antibody response in relative BAU of a 49-year-old female without pre-vaccine infection (Baseline) to a 75-year-old male with pre-vaccine infection (Covariates). Vertical black lines indicate vaccination events. (E) Comparison of the same individuals as in (D) but at a fixed time point at 700 days after first vaccination.

We then incorporated these candidate effects into the NLME model and performed refitting and model selection on a total of 19 different models (<u>S6 Fig</u>). In total, seven covariate–parameter interactions were retained in the final model (Fig 4B, <u>S1 Table</u>), as the corresponding covariate-effect parameters were significantly different from zero and improved the BICc by more than 10 (<u>S6 Fig</u>). Male sex affected both *r_1_* and *r_d_*, corresponding to weaker initial antibody production in the absence of pre-existing antibodies and slower antibody decline. Quantitatively, male sex was associated with an approximately 14.7% (95% CI: 9.6-19.4%) reduction in the effective production parameter *r_1_* and an approximately 7.1% (95% CI: 3.1-10.9%) reduction in the decay parameter *r_d_*. Age also affected both *r_1_* and *r_d_*: with increasing age, the effective production parameter decreased by about 1.9% (95% CI: 1.7-2.1%) per year, whereas the decay parameter decreased by about 0.6% (95% CI: 0.4-0.7%) per year, indicating a weaker but more prolonged response. Pre-vaccine infection had the strongest overall effect on the dynamics. First, it introduced non-zero initial antibody levels before the first vaccination; the corresponding average level was estimated at 5.3 BAU (95% CI, 3.3–8.4). Second, pre-vaccine infection altered both the production and decline parameters, reducing *r_1_* by approximately 85.6% (95% CI: 38.1-96.6%) and increasing *r_d_* by approximately 92.7% (95% CI: 83.5-102.4%), thereby shifting the overall response trajectory. Finally, cohort membership affected *A_0_*, with 641.4% (95% CI: 356.9-1103.1%) higher estimated initial antibody levels in Brescia than in Bologna, which would equal 34.0 BAU.

To assess the combined effects of these covariates and how they translate into substantial differences in predicted antibody trajectories, we consider the predicted antibody levels from the estimated population average and covariate effects parameters of a 49-year-old female individual without pre-vaccine infection (Baseline) and a 75-year-old male individual with pre-vaccine infection (Covariates) (Fig 4C-E). The predicted antibody levels differed strongly already after the first vaccination. These differences persisted over time, but slowly reduced and reached a turning point around six months after the third vaccination where the first individual surpassed the second in antibody levels. One year after the third vaccination the second individual would have approximately 27% lower antibody levels than the first individual. This comparison exemplifies how host covariates do not merely shift antibody levels at isolated time points, but act through distinct, mechanistically interpretable features of the response dynamics.

## Discussion

Antibody responses after COVID-19 vaccination vary widely between individuals, yet previous analyses have only incompletely resolved which covariates shape the dynamics of these responses in large heterogeneous populations. Here, we developed and calibrated a NLME model of vaccine-induced antibody dynamics using longitudinal data from more than 4,000 individuals. This complements previous smaller-scale dynamic modeling studies as well as larger observational studies that quantified associations with antibody levels but could not relate them to dynamic features of the response. Our analysis shows how age, sex, and prior infection differentially affect antibody production and waning dynamics after vaccination.

Overall, we identified seven statistically significant covariate–parameter relationships in the final model. The large cohort size enabled us to detect not only strong but also comparatively subtle covariate effects. In general, the findings were consistent with previous reports. For example, we found that male sex was associated with reduced initial antibody production, in line with Collatuzzo et al. [32], who reported lower antibody levels nine months after vaccination in men in a larger cohort of healthcare workers including those considered in this analysis. We also found that increasing age was associated with lower initial antibody production together with slower antibody decline. Taken together, these effects imply a weaker but more prolonged response in older individuals, consistent with observations reported by Leomanni et al. [11] in the same expanded cohort. The biological basis of this pattern remains uncertain, but it may reflect age-dependent changes in immune-cell activity, turnover, or persistence. While our model cannot resolve these mechanisms directly, it provides a quantitative framework for describing how age reshapes the temporal structure of the response. More broadly, this illustrates the added value of a dynamic modeling framework: covariates may be related to different components of the response in opposing ways, which cannot be resolved from cross-sectional studies alone.

Pre-vaccine infection had the strongest overall effect on the response dynamics. Beyond introducing non-zero initial antibody levels before first vaccination, it reduced the effective production and increased decline parameters of the model. A related phenomenon has been reported in other vaccination settings; for example, higher pre-existing antibody levels against meningitis were associated with a smaller fold-increase after vaccination [33]. However, we could not reconcile the faster antibody decline with previous studies. For example, Srivastava et al. [34] found individuals with pre-vaccine infection to have higher antibody levels up to 400 days post second vaccination compared to SARS-CoV-2 naive individuals and no difference after a third vaccination. In contrast, we found that antibody levels of an individual with pre-vaccine infection would be higher initially but could drop below that of one without pre-vaccine infection. One possible explanation is that pre-vaccine infection changes the balance between immediate antibody availability and the processes driving sustained response after vaccination. Future model extensions that include additional immune compartments, in particular memory-cell populations [35], may help disentangle these effects and clarify the mechanisms by which prior infection reshapes vaccine responses.

Our study has several limitations. First, the proposed model provides only a very coarse-grained description of a highly complex biological process. Vaccine-induced immunity involves multiple interacting cell types and regulatory pathways [20], many of which are not explicitly represented here. This was necessary as our in silico analyses showed that more detailed model formulations could not be estimated reliably under the data conditions available in this study, motivating the use of a simpler model. Indeed, the use of overly complex models yielded biased estimation results, highlighting the importance of in silico studies and the reporting of the respective results. Second, the available measurements were limited to longitudinal antibody titres and vaccination times, which restricts the degree to which estimated parameters can be interpreted mechanistically. Richer longitudinal immunological data, for example on memory B cells, plasma cells, or other immune compartments, would be required to support more detailed mechanistic models. Such data can in principle be collected, but cohort sizes are substantially smaller [36,37]. Third, the range of covariates considered was limited by the information available in the ORCHESTRA cohort. Additional variables, such as immunodeficiency, comorbidities, medication, or disease severity, may explain further heterogeneity and should be explored in future studies.

Finally, denser longitudinal sampling would reduce the need for filtering and would make it possible to account more explicitly for infections occurring during follow-up, rather than excluding affected individuals from the analysis.

Despite these limitations, our study provides an important new perspective and has several unique characteristics: First, it combines a large longitudinal cohort of more than 4,500 individuals with a mechanistic modeling framework, enabling the analysis of vaccine-induced antibody dynamics at a scale that exceeds most previous dynamic modeling studies. Second, the nonlinear mixed-effects approach allowed us to simultaneously characterize population-level response patterns and inter-individual variability, while directly linking host covariates to interpretable features of antibody production and decline. This provides information beyond that obtainable from cross-sectional analyses or regression models applied to isolated time points. Third, the availability of repeated measurements over an extended follow-up period enabled us to investigate how covariate effects accumulate over time and translate into long-term differences in antibody trajectories.

In conclusion, our modeling and analysis provide a dynamic view of the factors shaping antibody responses after COVID-19 vaccination. By linking host covariates to interpretable features of antibody production and decline, the study helps explain long-term heterogeneity in humoral immunity. These findings may inform future work on individualized vaccination schedules and strategies to modulate vaccine responses, including the use of adjuvants [15]. Moreover, the model could contribute to more comprehensive frameworks for assessing infection and reinfection risk [38–40]. In the future, coupling detailed models of antibody dynamics and infection risk with models of post-infection syndromes [41–43] could enable a more comprehensive assessment and prediction of short- and long-term disease burden while accounting for inter-individual variability.

## Author contributions

**Conceptualization**: Clemens Peiter, Jan Hasenauer, Paolo Boffetta

**Data curation**: Clemens Peiter, Emma Sala, Carlotta Zunarelli, Mahsa Abedini.

**Formal Analysis**: Clemens Peiter, Jan Hasenauer.

**Funding Acquisition**: Evelina Tacconelli, Paolo Boffetta, Jan Hasenauer.

**Methodology**: Clemens Peiter, Jan Hasenauer.

**Project Administration**: Michelle B. Hanson, Evelina Tacconelli, Paolo Boffetta, Jan Hasenauer.

**Resources**: Emma Sala, Giuseppe De Palma, Carlotta Zunarelli, Mahsa Abedini

**Supervision**: Jan Hasenauer.

**Visualization**: Clemens Peiter.

**Writing – Original Draft Preparation**: Clemens Peiter, Jan Hasenauer.

**Writing – Review & Editing**: All authors.

## Acknowledgements

The authors acknowledge the ORCHESTRA Working group. We thank all participants of the study for their contribution to this research.

## Supporting information

**S1 Text. Additional information on data curation, model simplification, and estimation bias for synthetic datasets with the original and simplified models.**

**S1 Table. Parameter estimates of the best start and lower and upper bound used to sample start parameters from a uniform distribution.** Values are rounded to the second decimal.

**S1 Fig. Analysis of parameter identifiability of the original model for synthetic data of a single individual.** (A) Model fit plot showing the best fit obtained after multistart optimization for a synthetic dataset created from the original model by dePillis et al. (B) Estimated parameter values for the synthetic dataset shown in blue. The grey dashed line indicates the true parameters and black dashed lines indicate parameter bounds used in the parameter estimation. (C) Profile likelihood plots for all four estimated parameters. (D-F) The subfigures show the same as in (A-C), but with a larger synthetic dataset.

**S2 Fig. Analysis of identifiability of our simplified model for synthetic data of a single individual.** (A) Model fit plot showing the best fit obtained after multistart optimization for a synthetic dataset created from the original model by dePillis et al. (B) Estimated parameter values for the synthetic dataset shown in blue. The grey dashed line indicates the true parameters and black dashed lines indicate parameter bounds used in the parameter estimation. (C) Profile likelihood plots for all four estimated parameters. (D-F) The subfigures show the same as in (A-C), but with a larger synthetic dataset.

**S3 Fig. Parameter estimates of synthetic datasets with the original and simplified models.** (A,B) Estimates of the original model. (A) Plot of the estimated likelihood showing the convergence in likelihood for our multistart approach for the ten best starts sorted by likelihood for each synthetic dataset. Each color represents a different synthetic dataset and lighter colors represent worse likelihoods. (B) Corresponding parameter estimates for the ten best starts for each of the three synthetic datasets. (C,D) Estimates of the simplified model.

**S4 Fig. Parameter estimates for the simplified model with all parameters subject to inter-individual variability.** (A) Plot of the estimated likelihood showing the convergence in likelihood for our multistart approach for the ten best starts sorted by likelihood. (B) Estimated parameter values for the ten best starts. (A,B) Lighter colors correspond to worse likelihood values.

**S5 Fig. Distribution of empirical and theoretical residuals for the best model fit.** (A) Comparison of empirical and theoretical cumulative distribution function of the individual weighted residuals (IWRES). (B) QQ-plot showing the theoretical and sampled quantiles of the IWRES.

**S6 Fig. Tested parameter-covariate relationships and associated BICc.** Based on the simplified model, linear parameter-covariate associations were included in the model and the parameters were re-estimated. Upset plot showing all tested models and the included parameter-covariate associations (e.g., an association between rd and sex is indicated by rd∼sex).combinations that were tried, are listed on the left in the format “parameter_name-covariate”. job_title_group separates healthcare workers into two groups: one with intense patient contact, the other without. prev_infection is present when an individual has experienced a pre-vaccine infection.

**S7 Fig. Convergence of parameter estimation for the original model.** (A) Plot of the estimated likelihood showing the convergence in likelihood for our multistart approach for the ten best starts sorted by likelihood. (B) Estimated parameter values for the ten best starts. (A,B) Lighter colors correspond to worse likelihood values.

