## Supplementary material for "Linking Host Covariates to COVID-19 Vaccine-Induced Antibody Dynamics in a Large Healthcare Worker Cohort": S1 Text

### Supporting information

#### Filtering steps applied to the dataset to arrive at the analysis set

Data were collected within the "Connecting European Cohorts to Increase Common and Effective Response to SARS-CoV-2 Pandemic" (ORCHESTRA) project and have been described previously by Leomanni et al. [1]. We were provided with data from two different ORCHESTRA study sites in Italy: Bologna and Brescia. From a dataset with initially 5,719 individuals, we first filtered out individuals with missing covariate information (n=1) and less than four measurements (n=1). Then, we filtered vaccine non-responders (n=48), i.e., individuals that had no measurable antibody response after two vaccinations. This group was too small to allow for a proper analysis. Finally, we filtered individuals that experienced an infection within the observation period (n=1,117). We define the observation period as the time between the first vaccination and the last measurement of anti-S antibodies or vaccination. Infections were either self-reported, identified by the presence of anti-nucleocapsid (N) antibodies, which is not expected for vaccination, or suspected by an increase of anti-S antibodies without the presence of a vaccination within a reasonable time frame before the measurement. To narrow down on suspected infections based on anti-S antibodies, individuals were selected based on several criteria: (i) anti-S antibody levels rise between two measurements and after 180 days after the last vaccination, (ii) the increase is larger than 25% of the previous antibody level (approximately equal to the measurement noise that was estimated), and (iii) the previous antibody level had been recorded within 180 days of the one in question. These selection criteria should prevent late vaccine responders and individuals whose antibodies appear to rise due to a measurement error from being excluded from the analysis. The selected individuals were excluded from further analysis, as the time of infection was unknown and could therefore not be modelled. Thus, we arrived at the analysis set of 4,553 individuals, with 761 from Bologna and 3,792 from Brescia.

#### Comparison of model from dePillis et al. (2023) and our model

We built our model on the model by dePillis et al. (2023) [2]. The authors' original model describes the antibody response to a vaccination where antibodies $A$ are produced in response to the presence of vaccine transfected cells $V$ (this is only a proxy for transfected cells, as their amount is unknown). The model is an ODE system and governed by the following equations:

$$dA/dt = r_1 V(t) + r_2 A(t)V(t) + A(t)\left(r_3 - r_4 A(t)\right)$$

$$dV/dt = u - k_1 V(t)/\left(k_2 + V(t)\right)$$

As most of the model has been described in the section *Material and methods* of the main manuscript, we will focus on the differences to the model used in this study. Critically, in the original model, the intrinsic antibody dynamics (in the absence of $V$) are modeled logistically. According to the authors, the original parameterization of the antibody decay as $r_4 A(t)^2$ represents the natural antibody decay and intra-species competition. The original model, however, showed poor convergence in parameter estimates for the experimental data at hand (S7 Fig) and biased parameter estimates for the synthetic data (S3 Fig A and B). Therefore, we replaced the logistic intrinsic dynamics with a simpler exponential decay term by removing the quadratic dependence on $A(t)$. Then, in the absence of transfected cells, that is for $V(t)=0$ , $dA/dt$ simplifies to

$$dA/dt = r_3 A(t) - r_4 A(t) = \left(r_3 - r_4\right) A(t) = r_d A(t).$$

Additionally, we restrict $r_d$ to be negative to prevent infinite exponential growth of antibodies. Thus, we arrive at our simplified model:

$$dA/dt = r_1 V(t) + r_2 A(t)V(t) - r_d A(t)$$

$$dV/dt = u - k_1 V(t)/\left(k_2 + V(t)\right).$$

### In silico testing of the modeling and parameter estimation pipeline

To test whether the parameter estimation workflow outlined in the section *Material and methods* of the main manuscript could reliably recover underlying parameters under conditions matching the real dataset, we performed a comprehensive assessment of the modeling and parameter estimation pipeline. Using the individual-level model and NLME framework, we obtained preliminary parameter estimates using the experimental data. Using these preliminary parameter estimates, we generated synthetic datasets with comparable population size, numbers of measurements per individual, sampling times, vaccination schedules and noise characteristics. In addition, we created synthetic datasets for a single individual with either four or eleven measurements. The larger dataset was created to check if more measurements could rescue a practical non-identifiability on an individual level. Compared to the experimental data, the synthetic measurements for a single individual were not censored.

These analyses showed that reliable estimation critically depended on the complexity of the individual-level model. In particular, when using the original model formulation considered as a starting point for our work [2], parameter recovery was not reliable under the study conditions considered here. Specifically, multistart optimization did not yield consistent convergence of parameter estimates across runs (S7 Fig). In addition, simulation experiments for a single individual showed that, with only four measurements, key parameters of the original model could not be identified reliably (S1 Fig). At the population level, this resulted in biased parameter recovery in the *in silico* setting (S3 Fig A and B). The bias was particularly pronounced for the ODE model parameters $\beta_{r_3}$ and $\omega_{r_3}$, as well as for the parameters $a_{Bol}$ and $a_{Bre}$, which constitute the absolute component of the mixed error model. Although the contribution of the absolute error component was small relative to the observed range of antibody levels and therefore of limited practical relevance, reliable recovery of the ODE model parameters was essential for the subsequent covariate analyses.

We therefore based the final analysis on the reduced model introduced above, in which the intrinsic antibody dynamics were simplified to first-order exponential decay. Repeating the *in silico* analyses with this reduced model showed accurate parameter recovery and robust estimation performance under the study conditions considered here (Fig 2, S2 Fig, S3 Fig C and D). These results provided an additional rationale for the final model choice and support the reliability of the subsequent analyses of the experimental data.
