## Supplementary material for "Linking Host Covariates to COVID-19 Vaccine-Induced Antibody Dynamics in a Large Healthcare Worker Cohort": S1 Table

| Parameter | Value | Standard error | Initialization lower bound | Initialization upper bound |
|---|---|---|---|---|
| **Mean** | | | | |
| $\beta_{A_0}$ | 1.66 | 0.23 | 0 | 3 |
| $\beta_{r_1}$ | 1.34 | 0.06 | -5 | 0 |
| $\beta_{r_d}$ | -5.44 | 0.04 | -8 | -3 |
| **Standard deviation** | | | | |
| $\omega_{A_0}$ | 1.25 | 0.06 | 3 | 3 |
| $\omega_{r_1}$ | 0.69 | 0.01 | 3 | 3 |
| $\omega_{r_d}$ | 0.38 | 0.01 | 3 | 3 |
| **Fixed across individuals** | | | | |
| $\beta_{r_2}$ | -0.47 | 0.02 | -5 | 0 |
| $\beta_{s_{Bre}}$ | 0.03 | 3.00E-3 | 0 | 2 |
| $a_{Bol}$ | 3.43 | 1.29 | 10 | 10 |
| $b_{Bol}$ | 0.28 | 6.07E-3 | 0.8 | 0.8 |
| $a_{Bre}$ | 9.79 | 1.79 | 10 | 10 |
| $b_{Bre}$ | 0.29 | 3.60E-3 | 0.8 | 0.8 |
| **Fixed value** | | | | |
| $k_1$ | 10 | – | – | – |
| $k_2$ | 50 | – | – | – |
| **Covariate effects** | | | | |
| $\gamma_{r_d}^{age}$ | -6.06E-3 | 8.45E-4 | 0 | 0 |

| | | | | |
|---|---|---|---|---|
| $\gamma_{r_1}^{age}$ | -0.02 | 1.17E-3 | 0 | 0 |
| $\gamma_{r_d}^{sex=male}$ | -0.07 | 0.02 | 0 | 0 |
| $\gamma_{r_d}^{inf.}$ | 0.66 | 0.02 | 0 | 0 |
| $\gamma_{r_d}^{sex=male}$ | -0.07 | 0.02 | 0 | 0 |
| $\gamma_{r_1}^{inf.}$ | -1.94 | 0.75 | 0 | 0 |
| $\gamma_{A_0}^{data=Brescia}$ | 2.00 | 0.25 | 0 | 0 |
