## Supplementary figures and images for "Linking Host Covariates to COVID-19 Vaccine-Induced Antibody Dynamics in a Large Healthcare Worker Cohort"

### S1 Fig

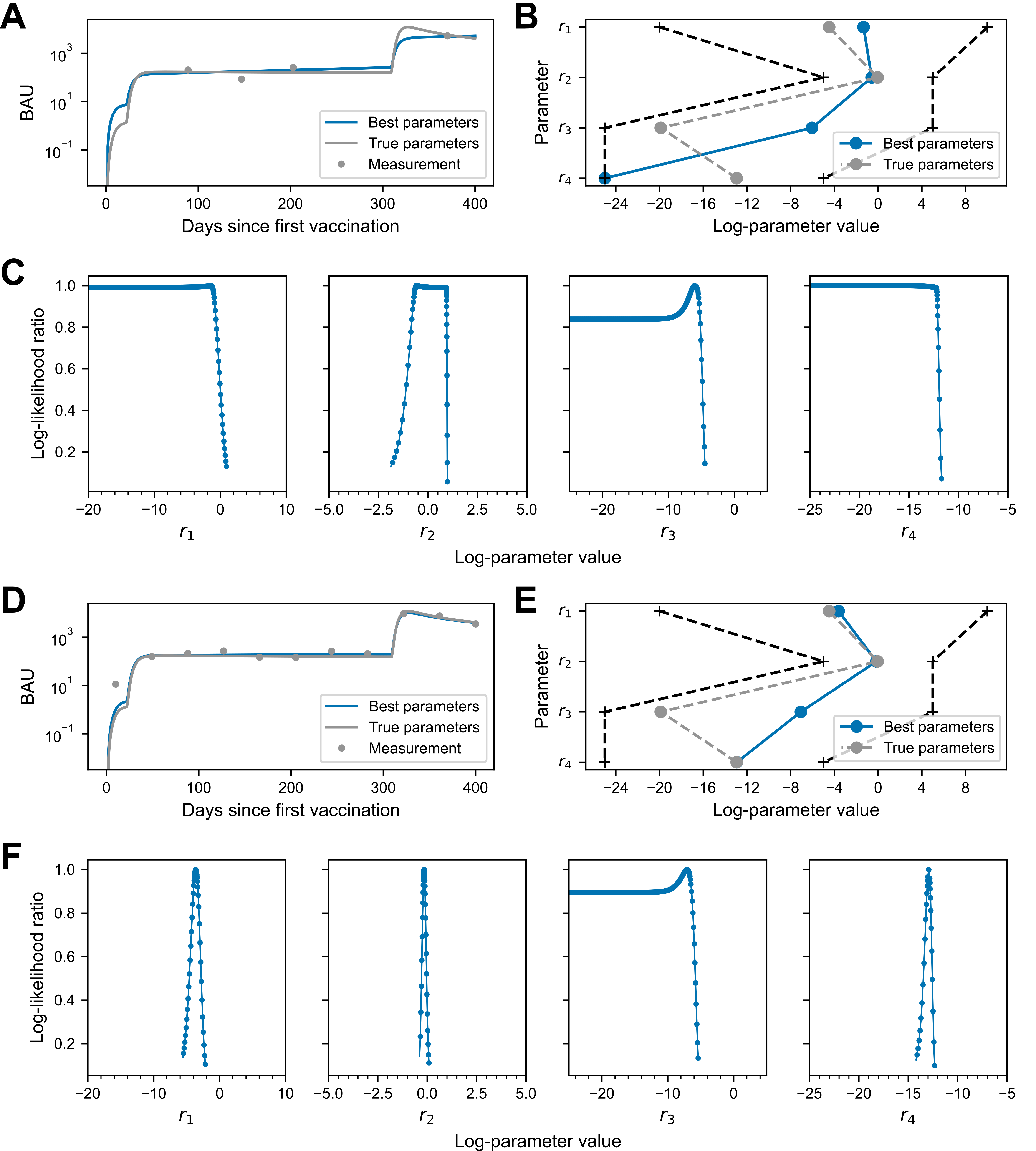

### S2 Fig

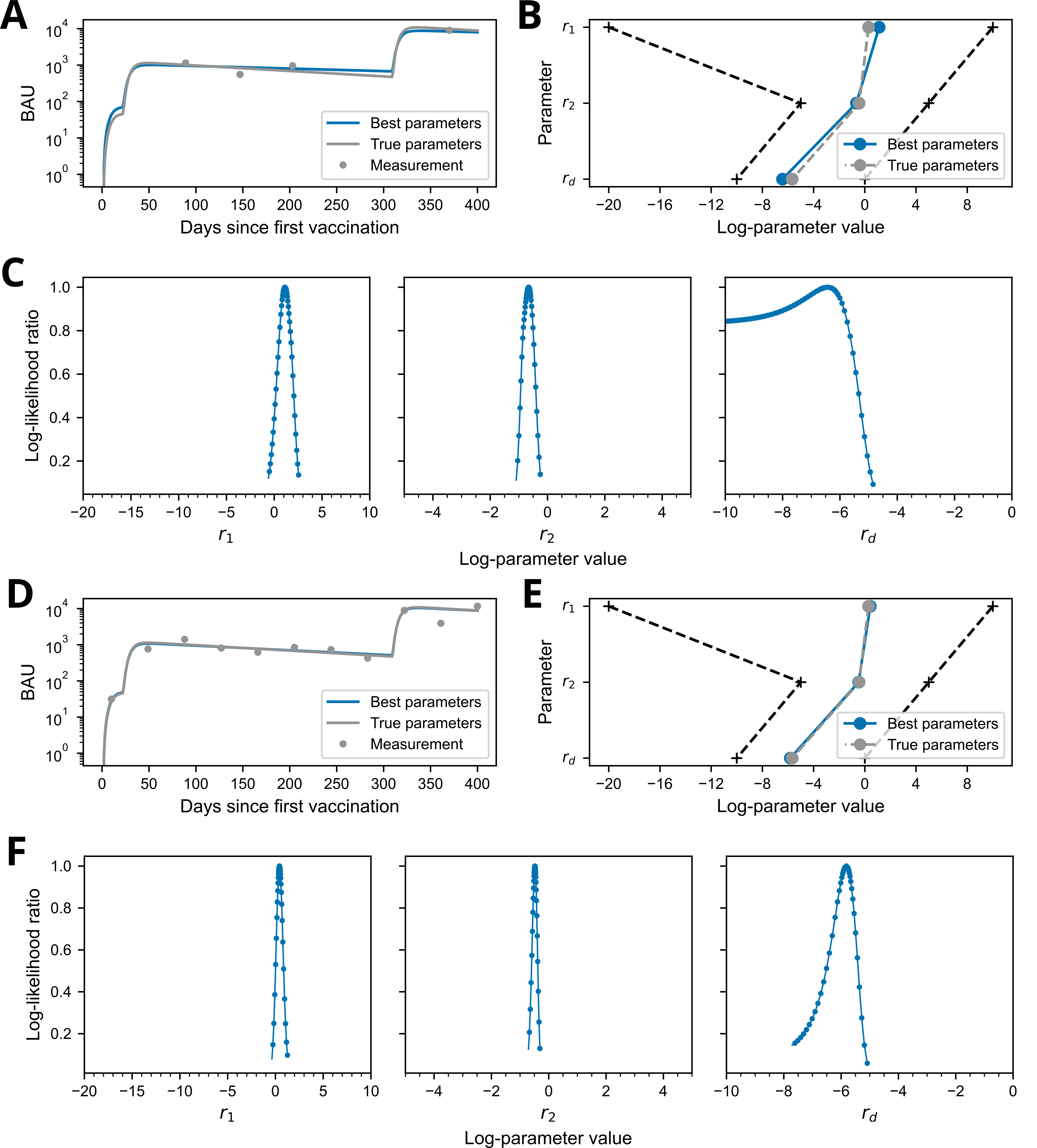

### S3 Fig

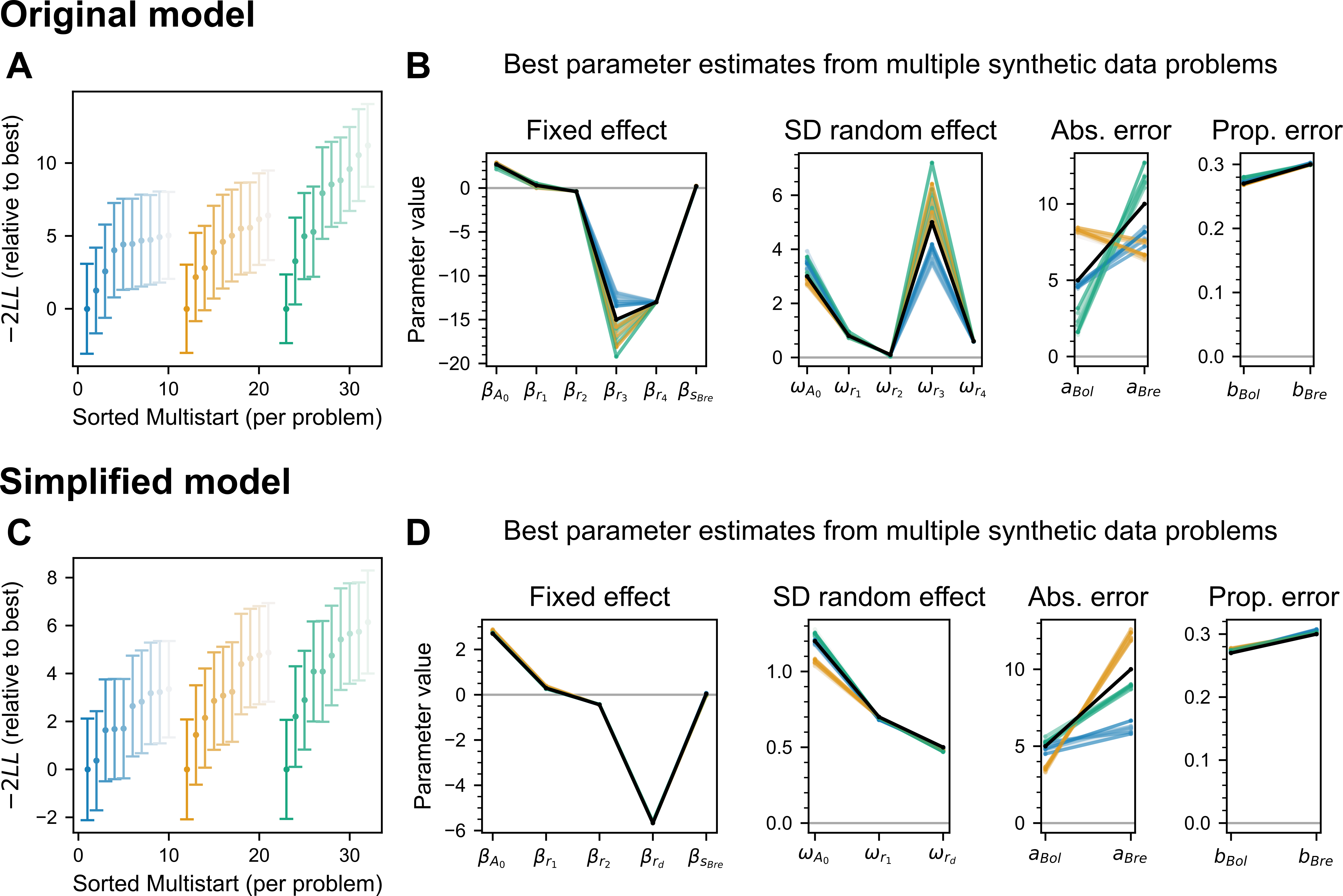

### S4 Fig

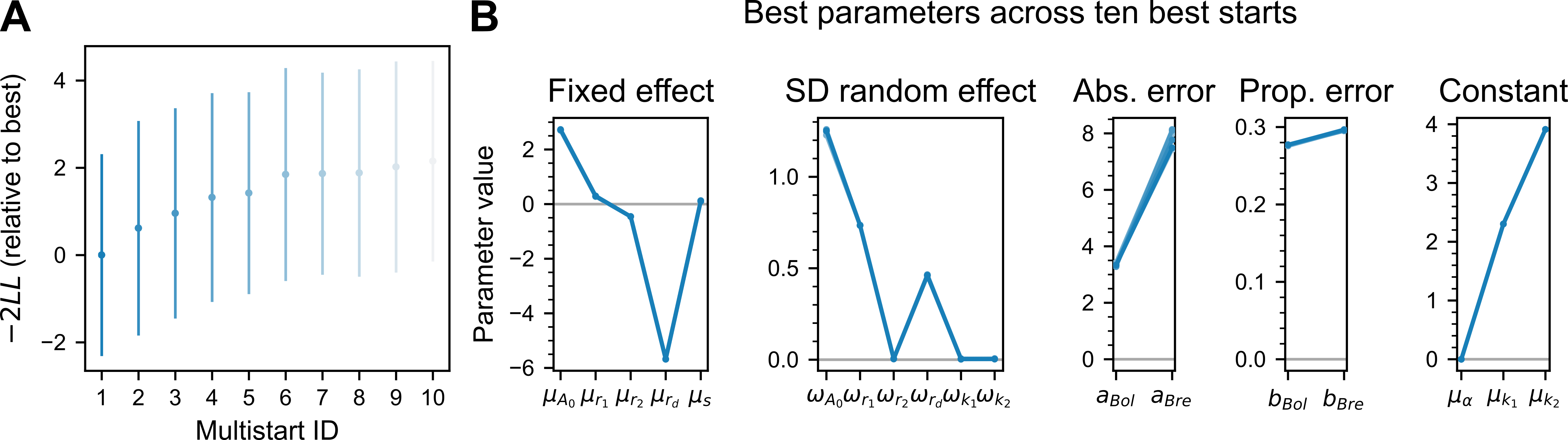

### S5 Fig

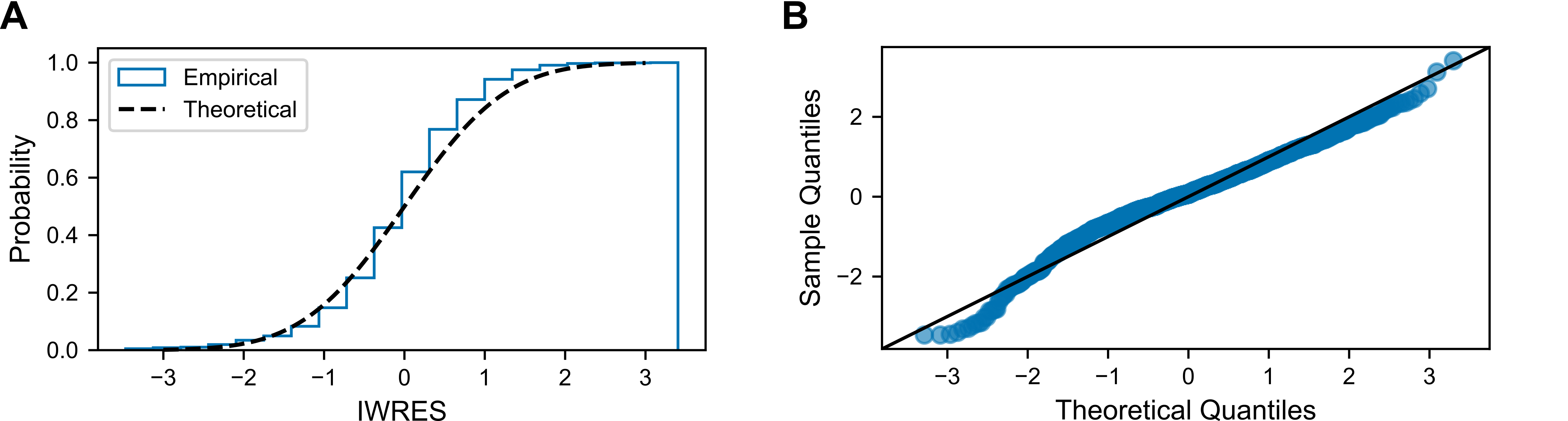

### S6 Fig

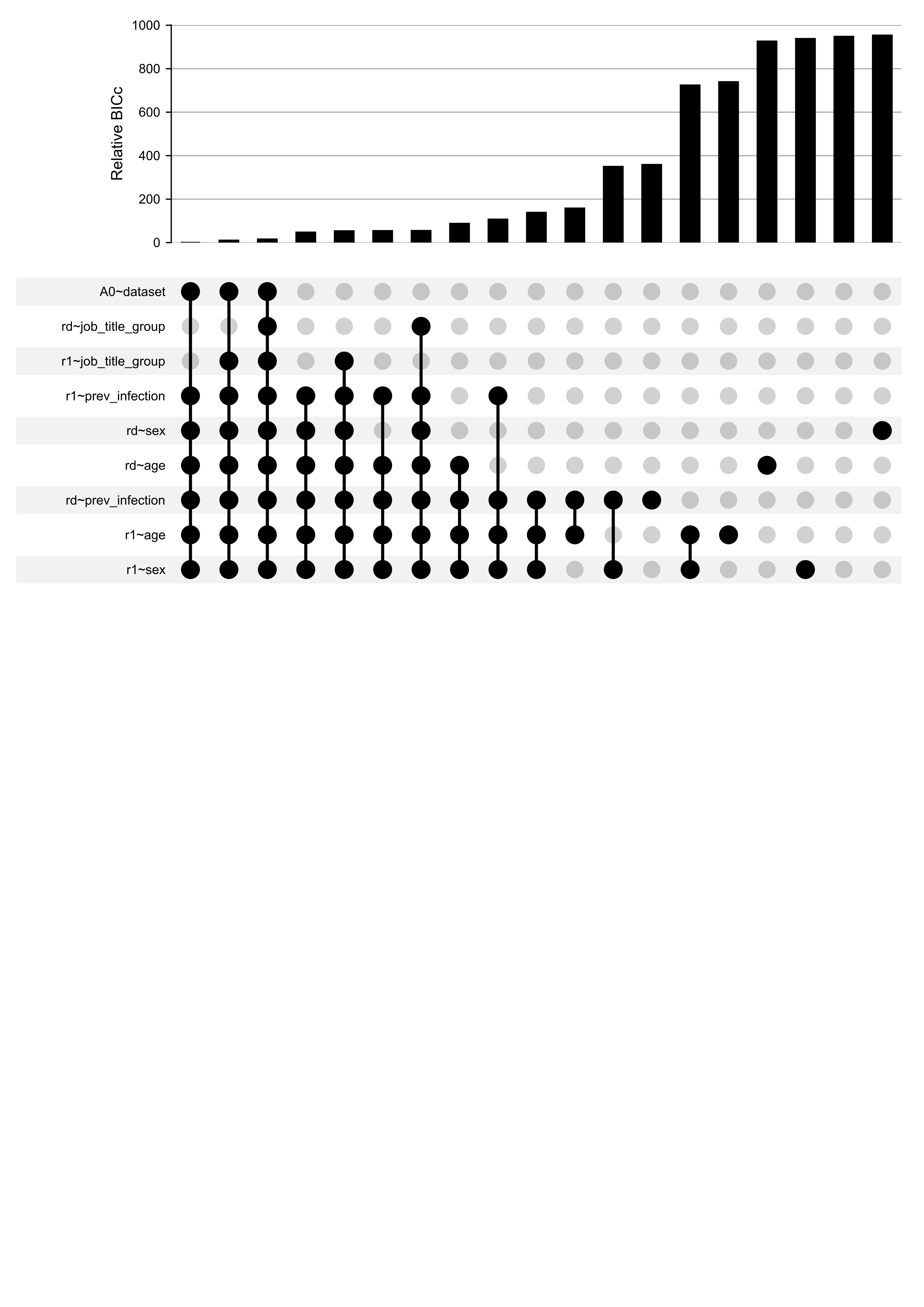

### S7 Fig

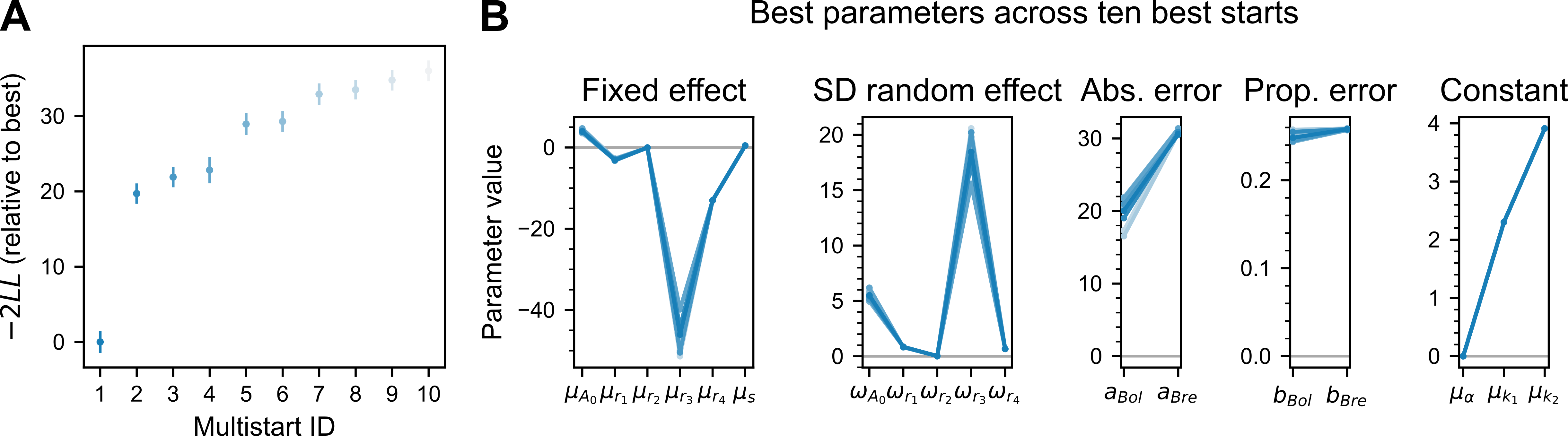
